# Differential maturation of mRGC pupillary constriction and circadian rhythmicity in typically developing human infants

**DOI:** 10.64898/2026.09.25.754365

**Authors:** Pär Nyström, Lilja Kristín Jónsdóttir

**Affiliations:** Department of Psychology, Uppsala University, Sweden

**Keywords:** Intrinsically photosensitive retinal ganglion cells, mRGC, pupillometry, circadian rhythms, sleep patterns, infants, development

## Abstract

Intrinsically light sensitive retinal ganglion cells through melanopsin photoreception (mRGCs) are important for fundamental regulatory brain processes such as circadian rhythms, mood, acute sleep, and pupillary constriction. While previous research in animal models indicates temporal differences in maturation of mRGC mediated projections, corresponding developmental patterns have not been studied in human infants. To elucidate the roles of mRGCs in early development of regulation of pupillary constriction and sleep/wake patterns, we here explore the maturation of mRGC mediated photoentrainment in human infants aged 1 to 12 months, using non-invasive eye tracking pupillometry and sleep diaries. Although the mRGCs relay rod/cone activation in a range of light intensities, it is possible to separate rod/cone/mRGC activation due to the photoreceptors’ color sensitivity and temporal responsiveness. We therefore used sustained red and blue light stimulation after partial dark adaptation to characterize developmental changes in wavelength-dependent mRGC-relayed pupillary responses. Infants’ pupil responses indicate a prolonged post-natal mRGC maturation to the shell of the olivary pretectal nucleus during the first year. This response was separate from infants’ sleep/wake patterns, which parallels findings from cellular research on murines: the mRGCs have separate pathways for photoentrainment of pupillary constriction and sleep/wake patterns, which develop with different temporal dynamics. Together, these findings provide the first characterization of developmental changes in mRGC-relayed pupillary responses in human infants, and an initial step towards bridging human mRGC network development with murine and human adult mRGC research.

**Statement of significance:** The retina contains rods, cones, and a more recently discovered photoreceptor called intrinsically photosensitive retinal ganglion cells (mRGC). One subtype of mRGCs project to the brain’s main area for circadian rhythms, the suprachiasmatic nucleus, and control photoentrainment of circadian rhythms such as day/night sleep patterns. While mRGCs have been extensively studied in murines and with adults there are no previous studies on typically developing infants. We here fill this gap and examine infant mRGC mediated activation using pupillary responses to red and blue light in relation to sleep/wake behavior, at an age range where light entrained sleep patterns emerge. The results parallels findings in animal research, and thereby start bridging literatures in animal research, adult sleep, and infant sleep.

## Introduction

Melanopsin containing retinal ganglion cells (mRGC), also called intrinsically photosensitive retinal ganglion cells (ipRGCs), is a third class of photoreceptor in the retina along with rods and cones, that play a key role in regulatory functions in both pupillary responses and circadian entrainment. Through their projections to subcortical structures involved in arousal and autonomic control, mRGCs are fundamental for the brain’s capacity to regulate arousal and environmental responsiveness. These regulatory functions develop early in life, and are essential aspects of healthy development. Adequate sleep is critical for typical brain development (Lokhandwala & Spencer, 2022), and in humans the adaptation to a day/night circadian rhythm is an important milestone for both the infant (and its parents). Regular sleeping patterns typically emerge around 8 weeks after birth (Rivkees, 2003), with progressively less daytime sleep during the first years (Lokhandwala & Spencer, 2022). While day/night circadian rhythms in adults have been found to be light entrained through mRGC mediated projections (Kawasaki & Kardon, 2007; Mure et al., 2019), the development of mRGC networks in the typically developing infant brain has not been studied previously.

The discovery of mRGCs at the turn of the millennium sparked new interest into the light sensitivity of the visual system. It was found that mRGCs are both *extrinsically* activated by rods and cones at low to medium light intensities (scotopic lighting < 0.01 cd/m^2^ and mesopic, 0.01 cd/m^2^ - 3 cd/m^2^) and *intrinsically* activated by melanopsin light absorption at medium to high light intensities (photopic lighting, > 3 cd/m^2^) (Altimus et al., 2010; Aranda & Schmidt, 2021; Güler et al., 2008; Schmidt et al., 2008, 2011). The melanopsin photopigment is encoded by the gene Opn4, which can be targeted by immunohistochemical and genetic tools to study the visual circuits of murines. Patch current-clamp recordings and multielectrode arrays recordings have mapped both the spectral sensitivities and temporal dynamics of the mRGCs (Aranda & Schmidt, 2021), and while it was initially thought that the mRGCs consisted of a single uniform cell population, it is now known that the mRGCs consists of several types, M1-M6 in murines, which project to different brain nuclei, have different response characteristics, and have different behavioral functions (Aranda & Schmidt, 2021; Mure et al., 2019; Schmidt et al., 2011; Schmidt & Kofuji, 2009).

The M1 cell type is most intrinsically sensitive to light, but does only marginally contribute to the image forming visual system, in contrast to M2-M6 (Aranda & Schmidt, 2021). Instead, the M1 type is the main player in the non-image forming visual system by projections to multiple brain areas: the suprachiasmatic nuclei (SCN) and intergeniculate leaflet (IGL) for circadian photoentrainment; the shell of the olivary pretectal nuclei (OPN) for pupillary light reflexes; perihabenular nucleus (pHb) for mood; ventrolateral preoptic area (VLPO) and lateral hypothalamic are (LH) for acute sleep; and multiple other hypothalamic regions (Schmidt et al., 2011; Schmidt & Kofuji, 2009). Notably, the projections to SCN and OPN are separated by distinct subpopulations of mRGCs (Brn3b-negative M1 and Brn3b-positive M1 respectively) (Chen et al., 2011), which opens up a theoretical separation between circadian entrainment and pupillary constriction also in humans.

Cellular research on mRGCs have typically been done using murine models, but recent translations to human mRGCs in vitro raise important differences. For example, only three mRGCs subtypes has been confirmed in humans compared to murines’ M1-M6, where two human types are suggested to correspond to mouse M1 and M2 (Mure et al., 2019). Also, humans are diurnal and have more cones than nocturnal mice and rats at a 20:1 ratio (McDougal & Gamlin, 2010; Mure, 2021; Mure et al., 2019), and the CRX gene which encodes the cone-rod homeobox transcription factor during development has important temporal expression differences (Bibb et al., 2001). There are also likely other developmental differences due to developmental allochrony (Matsuda et al., 2020), and although the development of mRGCs has been studied in rodents, showing temporal differences between postnatal innervation to SCN and OPN (McNeill et al., 2011; Schmidt et al., 2008; Tu et al., 2005), the developmental link between mRGC→SCN and mRGC→OPN pathways in human infants is unstudied.

To start filling this knowledge gap, we adapted a clinically targeted pupillometric method to measure rods, cones and mRGCs in adults (Kardon et al., 2009; Park et al., 2011) for use with infants. On a broader level we aimed to document an overview of the mRGC pupillary light response in typically developing infants, and more specifically, we tested the hypothesis that such mRGC mediated pupil responses are linked to the emergence of circadian sleep patterns. One widely used index of melanopsin-driven pupil activity is the post-illumination pupil response (PIPR), which quantifies the sustained pupil constriction that persists after stimulus offset following short-wavelength stimulation. Because the PIPR is considered a relatively specific marker of intrinsic melanopsin activity, it has become a standard outcome measure in chromatic pupillometry studies (Adhikari et al., 2015; Park et al., 2011). However, paradigms based on sustained illumination can also provide information about mRGC function during ongoing stimulation, reflecting the integrated contributions of rods, cones, and melanopsin-containing retinal ganglion cells. The present study builds on this literature by adapting a sustained-light infant paradigm intended to characterize developmental changes in mRGC-relayed pupillary responses.

## Methods

### Participants

79 typically developing infants cross-sectionally recruited between 1.6 and 12.0 months of age (35 girls, 36 boys, 8 unknowns due to missing questionnaires; grouped age statistics found in Supplementary table 1) and 10 adults (6 females, 4 males, 20-28 years, mean ± s.e.m. = 23.04 ± 0.76 years) participated in the study. Participants were stratified into four groups based on age: 1-4-month-olds (hereafter 2 months), 4-8-month-olds (hereafter 6 months), 8-12-month-olds (hereafter 10 months), and adults. Infant families were recruited from birth records or advertisements, and interviews with the parents ensured that the infants had no medical conditions or visual impairments relevant for the study. All adult participant and both parents of the infants signed informed consent before participation. The study was approved by a national ethics committee (Etikprövningsmyndigheten, Lund avdelning 2 medicin, dnr 2022-03706-01), preregistered at https://doi.org/10.17605/OSF.IO/9YRZN, and performed in agreement with national ethical regulations and the Helsinki Declaration.

### Design and procedure

All infants, parents and adult participants were dark adapted for approximately 5 minutes in a dim lit room with a controlled ambient light level of 0.01 lux while they were informed about the procedure of the study. We did not completely dark adapt the subject because we wanted to keep a clinically useful test paradigm similar to previous studies (Kardon et al., 2009), and complete dark adaption adds greatly to the procedure time. The participant was seated in front of a remote eye tracker (Tobii Spectrum, Tobii Technology, Danderyd, Sweden) with a sampling rate of 150 Hz. The stimulus was presented after a short eye tracker calibration sequence on the Spectrum computer monitor (531x296 mm) and consisted of a black screen (0.00 lux) baseline interval of 40 seconds, a 40 seconds red screen (RGB 1,0,0 giving 47 lux and a light spectrum peak at 619nm), and a 40 seconds blue screen (RGB 0,0,1 giving 47 lux and a light spectrum peak at 453nm). Four trials were presented in a red/blue/red/blue order, with baselines preceding, interleaving and appending the light stimulation. The 6-month- and 10-month-olds participated in a study with a larger test battery containing stimuli unrelated to mRGCs (video clips of faces and human interaction), which required shortened presentation times to 30 seconds light stimuli and 10 seconds baselines. To keep the attention of the infants, a small monochromatic video animation was presented at the center of the screen (176x100 mm) with low intensity difference to the background. Considering the absence of mRGC in the fovea (Mure, 2021), the pupillary impact of the video animation was considered a minor problem compared to eventual infant inattention and missing data when not using videos. All stimuli are publicly available on the Open Science Framework (https://osf.io/u42e3/). Note that thepresent design does not selectively isolate intrinsic melanopsin activation. Rather, it was developed as an infant-compatible assay of wavelength-dependent pupillary responses that are relayed through the mRGC system. At the irradiance levels used here, both rod/cone inputs and intrinsic melanopsin activity are expected to contribute to the response. Consequently, the present study cannot determine the relative contribution of intrinsic versus extrinsic ipRGC activation.

### Data preprocessing

All analyses were implemented in the Time Studio framework (Nyström et al., 2015) and are publicly available on the Open Science Framework (https://osf.io/u42e3/). The workflow includes both analysis scripts and raw datasets. Pupil diameter changes between samples exceeding 0.05mm were considered biologically implausible and such samples were excluded from the data. The eye tracker occasionally cannot track the pupil, resulting in missing data, and pupil traces shorter than 4 samples were considered unreliable and excluded. Missing data with gaps shorter than 3 samples were linearly interpolated. Data was resampled to 100Hz to get uniform time intervals between samples, and gaps in the pupil trace was extended by 3 samples both before and after the gap to remove artifacts from eye blinks (the estimated pupil size becomes smaller during eye lid closure). Perturbated trial means for each sample were calculated from randomly selected trial sample values (N = 1000) from a temporally weighted normal distribution with a standard deviation of 10 samples, to create a representative pupil trace devoid of artifacts from missing data (i.e. to fill in eye blinks and short gaps from flickering eye tracking), see Figure 1a.Dependent variables. The transient relative constriction was calculated as in our previous studies (Nyström et al., 2018): A^02^ − A^m2^)/A^02^, where A^0^ is the average pupil diameter-5 to 0 seconds before onset of light stimulation, and A^m^ is the minimum pupil diameter in the interval 0 to 3 seconds relative stimulus onset.

**Figure 1.**
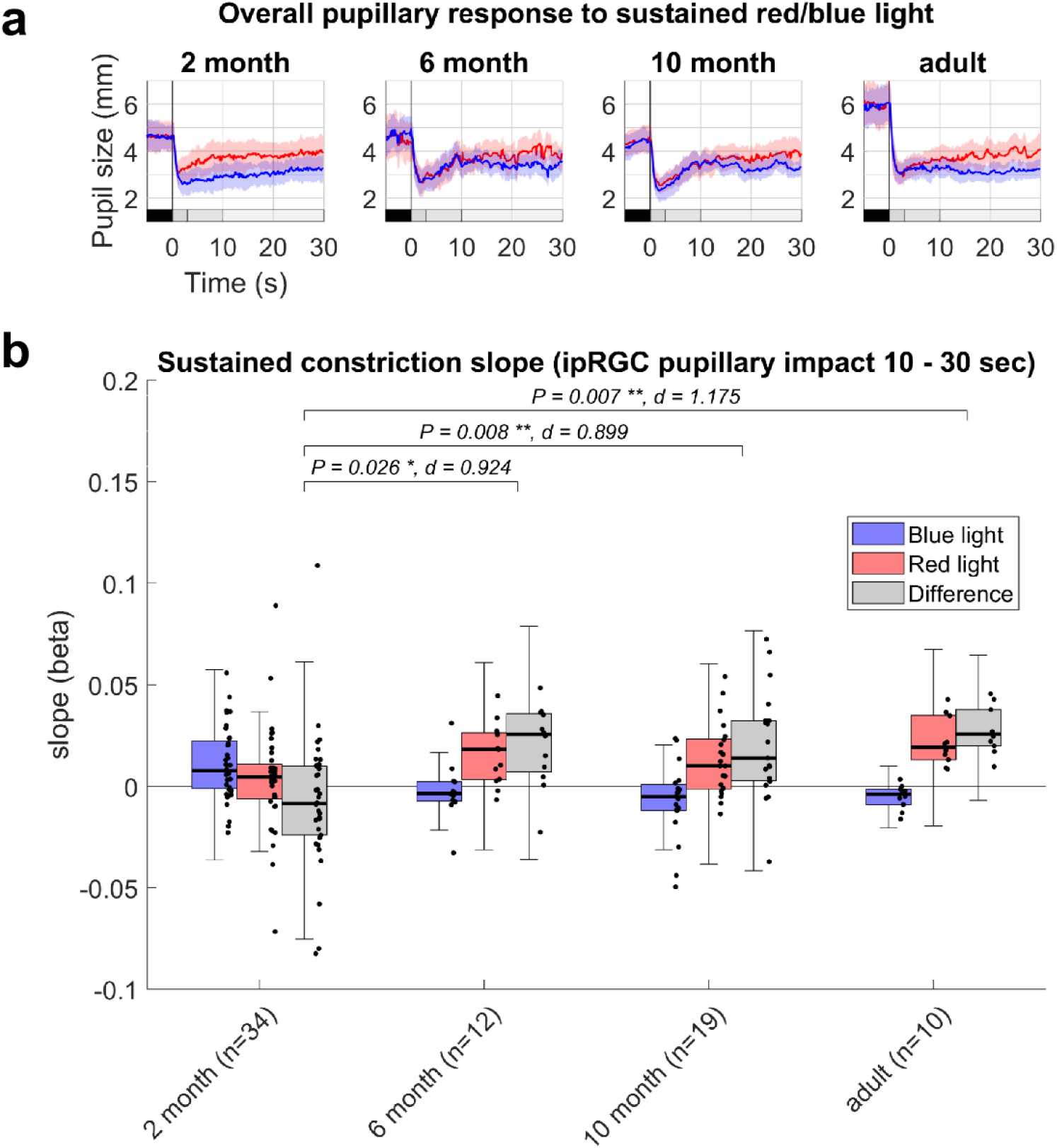
Data overview and developmental differences. (**a**) Panels show raw pupil sizes for all age groups. Lines represent perturbated trial means (see Methods), and shaded areas show standard deviations. Grey bars at bottom show the different analysis intervals: black is baseline-5 – 0 seconds relative light onset, dark grey is transient interval at 0 – 3 seconds where differences between red and blue relative constriction change between ages, F^3, 59^ = 10.58, P < 0.001; medium grey is escape interval 3 – 10 seconds where the difference between red and blue slopes of the pupil traces (Δβ) change between ages, F^3, 68^ = 5.33, P = 0.002; and light grey indicates our main analysis that shows the sustained interval 10 – 30 seconds where Δβ decrease between ages, F^3, 71^ = 7.28, P < 0.001. (**b**) Post-hoc test of developmental differences in sustained Δβ shows differences between the 2-month-olds and the other groups, with increasing differences between red and blue light indicating developmental changes in sustained mRGC-relayed pupillary responses. Only the difference values, Δβ, which we use to index mRGC mediated activation, were used for statistical analysis: blue and red is provided to illustrate data variability and integrity. We found no artifacts or noise in measurements that could explain outlier values, so all values were included in the analyses (see supplementary materials for inspection of all data as figures or in the TimeStudio (Nyström et al., 2015) scientific workflow). However, exploratory analyses without outliers showed larger effects between all groups, with P < 0.001 for all tests. Effect sizes are Cohen’s d, which are presented together with P-values above significance brackets. Center line is median; box limits are upper and lower quartiles; whiskers are 1.5x interquartile range; points are individual participant values.

The rationale of using sustained bright light stimulation is based on mRGCs role in maintaining sustained pupillary constriction, which is a summation of the adapted rod/cone response and the steady-state mRGC activation. To capture the adaption of the pupillary responses, which cannot be estimated with averaged constriction measures, we fit the slope of pupil diameter over time (*β*). For the calculation of pupillary escape (3 to 10 seconds) and sustained constriction (10 to 30 seconds), these intervals of interest was split into two parts with equal duration. Trials with less than 10% valid data in any part were excluded to ensure that slopes (*β*) calculated by linear regressions represented the whole interval well. Pupillary escape looking times in all trials (n = 271) were 85.2% ± 0.1 (s.e.m.) and sustained interval looking times (n = 270) was 76.1% ± 0.1 (s.e.m.). All trial slopes were visually inspected (data available at https://osf.io/u42e3/), but no trials were rejected because of implausible data. The difference in slopes between blue and red light stimulation is hereafter denoted Δ, and “sustained Δ*β”* was our primary measure for mRGC mediation activation, as stated in our preregistration (see Methods).

Previous literature has also used the post-illumination pupillary response (PIPR) as a marker of mRGC activity. Although the PIPR is frequently used as a marker of melanopsin activity, the present experiment was optimized for sustained pupillary recordings in awake infants rather than for PIPR estimation. Consequently, our primary outcome was the difference in sustained constriction slopes during ongoing stimulation. This measure should therefore be interpreted as reflecting wavelength - dependent mRGC-relayed pupillary activity rather than a direct measure of intrinsic melanopsin function.

### Behavioral data

All parents filled in the Brief Infant Sleep Questionnaire (BISQ) online in their homes after the eye tracking recording session. The parents of the 2-month-olds also filled in a sleep diary for 7 days and nights in their homes using pen and paper. The sleep diary was transcribed to digital form with 15-minute precision sleep/wake intervals, and sleep diary parameters were extracted as presented in Supplementary table 1.

### Statistical analysis

All statistical analyses were two-sided with an alpha of 0.05. We used 1-way ANOVAs to test overall group pupil constriction differences between infant groups and two sample *t* tests for *post-hoc* comparisons (using Bonferroni correction for families of 3 tests: 2-month vs 6-month, 2-month vs 10-month, and 6-month vs 10-month; the adult group was excluded because it was considered demonstrational and would inflate the statistical effects of developmental change). For Δ*β* analyses in relation to sleep diary measures, we performed linear regressions using the Matlab fitlm function. Q-Q plots and visual inspection of residuals were used to assess the assumptions and viability of statistical test, and all tests were found appropriate.

In the preregistration of this study we proposed linear mixed models for all statistical analyses, using the MATLAB fitlme function and with age as a continuous variable. These complementary analyses are presented in Supplementary Table 2 and gives conceptually identical results.

## Results

Our results show significant age-related developmental effects in three different measures motivated by previous research (Kardon et al., 2009; Kawasaki & Kardon, 2007; McDougal & Gamlin, 2010; Park et al., 2011): 1) the transient relative pupil constriction directly after light onset show less blue-red differences between 2-month-olds and 10-month-olds (Supplementary Figure 1), 2) the slope of pupillary escape between 2.5 – 10 seconds after light onset show less blue-red differences between 2-month-olds and 10-month-olds (Supplementary Figure 2), and 3) the slope of the sustained pupillary constriction between 10-30 seconds after light onset shows increased blue-red differences for 6-month-olds and 10-month-olds in comparison to 2-month-olds (Figure 1b, statistics found in figure and figure text). Overall, the pattern of effects suggests that mRGCs pupillary integration develop long after birth in humans, which parallels previous findings in mice (McNeill et al., 2011; Tu et al., 2005).

To investigate the relation between mRGC mediated activation (sustained Δ*β*) and the developing circadian rhythms and sleep/wake patterns, we let parents of the 2-month-olds track their infant’s sleep using sleep diaries for seven days and nights (Figure 2a) and fill in the Brief Infant Sleep Questionnaire (BISQ). Descriptive statistics for all measures are found in Supplementary Table 1. Statistical tests showed no significant effects between the sustained Δ*β* and any of the sleep measures (Figure 2b, statistics found in subplot titles), and follow up Bayesian regressions supported the null hypothesis with BF^01^ Mean = 2.57 ± 0.15 s.e.m. (median night-day sleep duration BF^01^ = 2.514), using the JASP 0.17.1.0 statistical software with default parameters.

**Figure 2.**
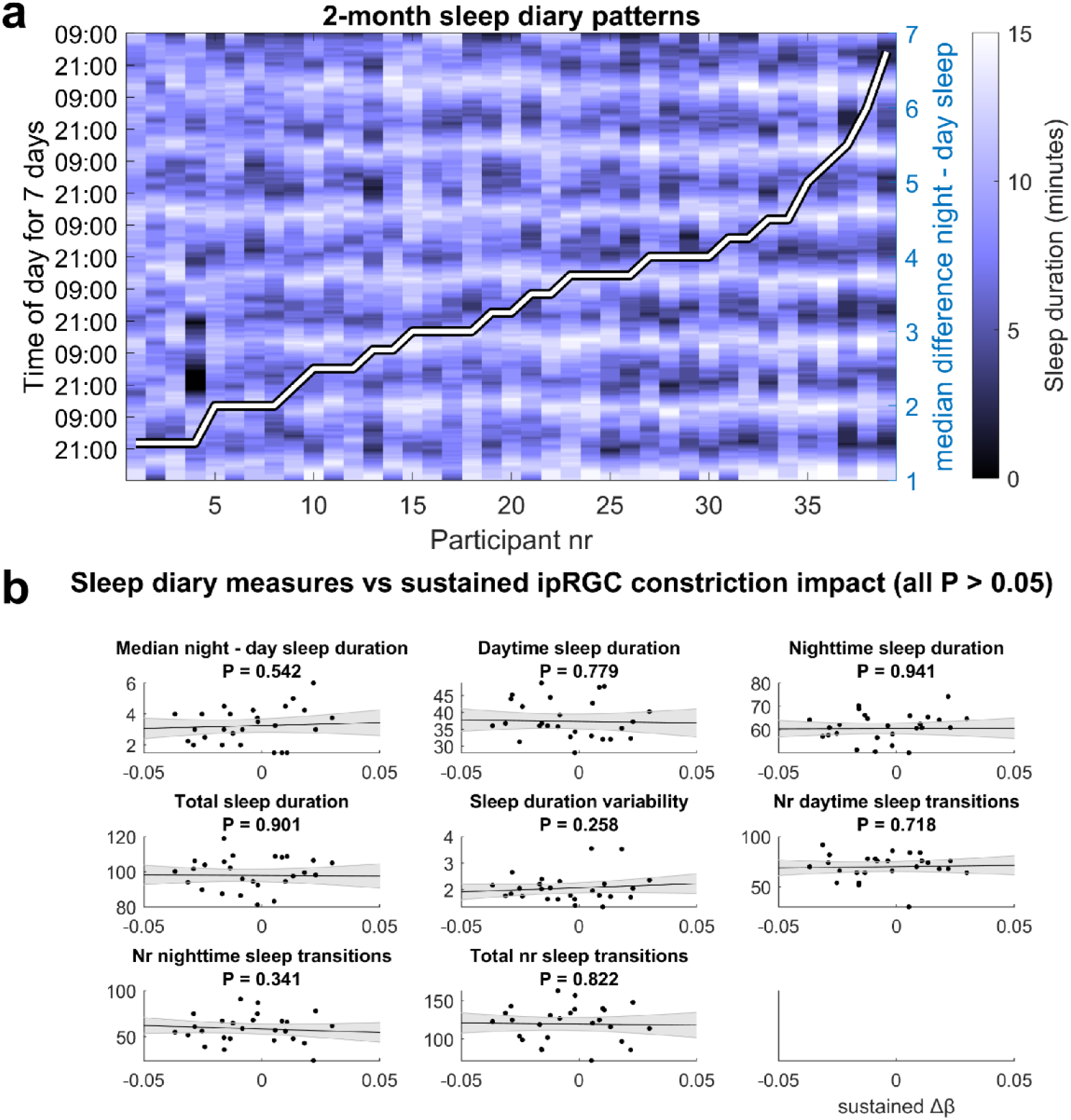
Sleep diary transcriptions and relation to sustained Δβ. (**a**) Background color image show parent scored sleep diaries, coded with 15-minute precision and smoothed vertically for demonstrational purposes. Participants are sorted horizontally based on our main measure of adaption to circadian sleep/wake rhythms: the median difference between night and day sleep duration, which is shown as a white line and that ranges between 1.5 and 6.75 hours. Note how a horizontal stripe pattern appear from left to right, showing the emergence of day/night rhythmicity in our sample (**b**) Subpanels show the relation between sustained Δβ (x-axes) and our sleep diary measures (y-axes). None of these analyses were statistically significant, suggesting decoupling of mRGC→SCN and mRGC→OPN shell development.

## Discussion

We here tested the hypothesis that pupil responses are linked to the emergence of circadian sleep patterns, through common mRGC mediated activation, and found support for the null hypothesis. Although this was not in line with our preregistered hypothesis, it is in line with discrepancies between developmental trajectories of mRGC→OPN shell innervation and mRGC→SCN innervation in murines (McNeill et al., 2011; Tu et al., 2005), which enables mRGC pupillary constriction to precede the emergence of light entrainment of circadian rhythms. Although we show a gradual maturation of the mRGC mediated pupillary constriction network from 2 to 12 months of age, it is important to emphasize that this system is by no means inactive before then. The rapid rod+cone pupillary light reflex is relayed through mRGCs (Aranda & Schmidt, 2021; Mure et al., 2019), and light evoked pupillary constriction has been shown already in premature infants after week 33 (Hanita et al., 2009; Ikeda et al., 2015; Robinson & Fielder, 1990). The expression of opsin genes starts even earlier, and mRGC melanopsin is already produced by 8-9 weeks post-conception (Tarttelin et al., 2003). The functional role of this early expression (or the delayed expression of rod and cone opsin genes) is still unclear since melanopsin expression long predates the development of the optic nerve (Bibb et al., 2001; Tarttelin et al., 2003). However, the early expression of melanopsin and the prolonged maturation of mRGC networks are conserved between species (Tarttelin et al., 2003), indicating an evolutionary robust mechanism likely to be crucial for typical brain developmental.

The current study has several limitations worth mentioning. First, recording pupillometric data in infant populations using stimuli with long duration is challenging due to infants’ limited attention. Although infants blink far less than adults, they have different eye shapes that can make the eye tracker provide noisy or missing data. Together with the additional missing data from head turns and other movements, the signal is not as solid as adult data. We therefore visually inspected all pupil traces and included figures of all pupil traces in the Supplementary Material. Further, we performed exploratory analyses with outliers removed, and still found the main results valid, which suggests that data quality was sufficient to answer our research question.

Second, related to the issue of attention, we used a regular computer monitor to present an attention capturing animated videoclip as stimuli, instead of stationary monochromatic light. This was a deliberate choice after piloting adults with both screen animations and monochromatic light, where we found similar differences in *Δβ* for both types of stimuli. While we cannot completely rule out that pupillary responses differ between stimuli types, the feasibility of completing recordings without infants fussing or otherwise cancelling the session favored the screen based approach.

Third, we only assessed sleep using parental reported sleep diaries, and did not use actigraphs or polysomnography to obtain more objective measures of sleep and activity levels. We can only speculate how these different methodological approaches would differ, and we therefore leave the question whether objective sleep measures would show statistically significant links to pupillometric measures for future studies.

While our findings are the first to document development of the mRGC mediated pupillary constriction in typically developing infants, we believe this to only be the first step in a larger endeavor. We show distinct maturation between 2 and 6 months of age, and that our measure of mRGC activation was not coupled with circadian sleep/wake rhythms, suggesting a functional separation between mRGC→SCN and mRGC→OPN pathways. This bears methodological considerations to important future questions, such as mRGCs role in infants’ acute versus circadian sleep from infancy to adolescence; seasonal effects on brain development of daylight exposure at different latitudes; or mRGCs roles in early sleep and later neuropsychiatric disorders, to name a few. Clearly, the practical limitations when studying typically developing infants poses difficult challenges while bridging the knowledge gap between cellular mRGC characteristics in murines and mRGC related behavioral characteristics in humans. Despite this, like others (Kardon et al., 2009; McDougal & Gamlin, 2010; Park et al., 2011), we believe that human pupillary responses to light provide a powerful model of the light sensitivity of human mRGCs. We also believe that this model is useful in infant research, and that developmental mRGC studies will be an important tool in the continued step-by-step pursuit of understanding the human brain and mind.

## Acknowledgements

The author is grateful to prof. G. Gredebäck for academic autonomy, and L. Hazer, E. Bodin, E. Hursti, M. Bravo, H. Nyström, E. Fredriksson, I. Cherco, A. Månson, O. Yttermo, H. Johansson, M. Jaghammar, and C. Beanier for data collection. This study was funded by G. Gredebäck’s KAW 2017.0284, and P. Nyström’s Swedish Research Council 2023-01503.

## Author contributions

P.N. designed the experiment, analyzed the data, and wrote the first draft of the paper. L.J. contributed with conceptual design, disseminated and discussed the results at ESCAP 2025 congress (symposium 19044), and critically revised the manuscript.

## Data availability

All data and the scientific workflow used to analyze the data is available at the Open Science Framework as a Timestudio project: https://osf.io/u42e3/

## References

Altimus, C. M., Güler, A. D., Alam, N. M., Arman, A. C., Prusky, G. T., Sampath, A. P., & Hattar, S. (2010). Rod photoreceptors drive circadian photoentrainment across a wide range of light intensities. Nature Neuroscience, 13 (9), 1107–1112.

Aranda, M. L., & Schmidt, T. M. (2021). Diversity of intrinsically photosensitive retinal ganglion cells: Circuits and functions. Cellular and Molecular Life Sciences, 78, 889–907.

Bibb, L. C., Holt, J. K., Tarttelin, E. E., Hodges, M. D., Gregory-Evans, K., Rutherford, A., Lucas, R. J., Sowden, J. C., & Gregory-Evans, C. Y. (2001). Temporal and spatial expression patterns of the CRX transcription factor and its downstream targets. Critical differences during human and mouse eye development. Human Molecular Genetics, 10 (15), 1571–1579.

Chen, S.-K., Badea, T. C., & Hattar, S. (2011). Photoentrainment and pupillary light reflex are mediated by distinct populations of ipRGCs. Nature, 476 (7358), 92–95.

Güler, A. D., Ecker, J. L., Lall, G. S., Haq, S., Altimus, C. M., Liao, H.-W., Barnard, A. R., Cahill, H., Badea, T. C., Zhao, H., & others. (2008). Melanopsin cells are the principal conduits for rod–cone input to non-image-forming vision. Nature, 453 (7191), 102–105.

Hanita, T., Ohta, H., Matsuda, T., & Miyazawa, H. (2009). Monitoring preterm infants’ vision development with light-only melanopsin is functional. The Journal of Pediatrics, 155 (4), 596–596.

Ikeda, T., Ishikawa, H., Shimizu, K., Asakawa, K., & Goseki, T. (2015). Pupillary size and light reflex in premature infants. Neuro-Ophthalmology, 39 (4), 175–178.

Kardon, R., Anderson, S. C., Damarjian, T. G., Grace, E. M., Stone, E., & Kawasaki, A. (2009). Chromatic pupil responses: Preferential activation of the melanopsin-mediated versus outer photoreceptor-mediated pupil light reflex. Ophthalmology, 116 (8), 1564–1573.

Kawasaki, A., & Kardon, R. H. (2007). Intrinsically photosensitive retinal ganglion cells. Journal of Neuro-Ophthalmology, 27 (3), 195–204.

Lokhandwala, S., & Spencer, R. M. (2022). Relations between sleep patterns early in life and brain development: A review. Developmental Cognitive Neuroscience, 101130.

Matsuda, M., Hayashi, H., Garcia-Ojalvo, J., Yoshioka-Kobayashi, K., Kageyama, R., Yamanaka, Y., Ikeya, M., Toguchida, J., Alev, C., & Ebisuya, M. (2020). Species-specific segmentation clock periods are due to differential biochemical reaction speeds. Science, 369 (6510), 1450–1455.

McDougal, D. H., & Gamlin, P. D. (2010). The influence of intrinsically-photosensitive retinal ganglion cells on the spectral sensitivity and response dynamics of the human pupillary light reflex. Vision Research, 50 (1), 72–87.

McNeill, D. S., Sheely, C. J., Ecker, J. L., Badea, T. C., Morhardt, D., Guido, W., & Hattar, S. (2011). Development of melanopsin-based irradiance detecting circuitry. Neural Development, 6, 1–10.

Mure, L. S. (2021). Intrinsically photosensitive retinal ganglion cells of the human retina. Frontiers in Neurology, 12, 636330.

Mure, L. S., Vinberg, F., Hanneken, A., & Panda, S. (2019). Functional diversity of human intrinsically photosensitive retinal ganglion cells. Science, 366 (6470), 1251–1255.

Nyström, P., Falck-Ytter, T., & Gredebäck, G. (2015). The TimeStudio Project: An open source scientific workflow system for the behavioral and brain sciences. Behavior Research Methods, 1–11.

Nyström, P., Gliga, T., Jobs, E. N., Gredebäck, G., Charman, T., Johnson, M. H., Bölte, S., & Falck-Ytter, T. (2018). Enhanced pupillary light reflex in infancy is associated with autism diagnosis in toddlerhood. Nature Communications, 9 (1), 1–5.

Park, J. C., Moura, A. L., Raza, A. S., Rhee, D. W., Kardon, R. H., & Hood, D. C. (2011). Toward a clinical protocol for assessing rod, cone, and melanopsin contributions to the human pupil response. Investigative Ophthalmology & Visual Science, 52 (9), 6624–6635.

Rivkees, S. A. (2003). Developing circadian rhythmicity in infants. Pediatrics, 112 (2), 373–381.

Robinson, J., & Fielder, A. (1990). Pupillary diameter and reaction to light in preterm neonates. Archives of Disease in Childhood, 65 (1 Spec No), 35–38.

Schmidt, T. M., Chen, S.-K., & Hattar, S. (2011). Intrinsically photosensitive retinal ganglion cells: Many subtypes, diverse functions. Trends in Neurosciences, 34 (11), 572–580.

Schmidt, T. M., & Kofuji, P. (2009). Functional and morphological differences among intrinsically photosensitive retinal ganglion cells. Journal of Neuroscience, 29 (2), 476–482.

Schmidt, T. M., Taniguchi, K., & Kofuji, P. (2008). Intrinsic and extrinsic light responses in melanopsin-expressing ganglion cells during mouse development. Journal of Neurophysiology, 100 (1), 371–384.

Tarttelin, E. E., Bellingham, J., Bibb, L. C., Foster, R. G., Hankins, M. W., Gregory-Evans, K., Gregory-Evans, C. Y., Wells, D. J., & Lucas, R. J. (2003). Expression of opsin genes early in ocular development of humans and mice. Experimental Eye Research, 76 (3), 393–396.

Tu, D. C., Zhang, D., Demas, J., Slutsky, E. B., Provencio, I., Holy, T. E., & Van Gelder, R. N. (2005). Physiologic diversity and development of intrinsically photosensitive retinal ganglion cells. Neuron, 48 (6), 987–999.

