## Supplementary materials for "Differential maturation of mRGC pupillary constriction and circadian rhythmicity in typically developing human infants"

Pär Nyström, 2026

**Supplementary Tables**

**Supplementary Table 1.** Descriptive data and infant group comparisons.

| Measure | 2 month | 6 month | 10 month | ANOVA |
|---|---|---|---|---|
| Age in months | N = 40<br>M = 2.07 ± 0.39 | N = 13<br>M = 6.74 ± 0.75 | N = 18<br>M = 11.00 ± 0.99 | $F_{2, 68}$ = 1185.51<br>*P* = 0.000 |
| Transient relative constriction | N = 25<br>M = 17.83 ± 10.14 | N = 10<br>M = 15.51 ± 12.22 | N = 18<br>M = 6.63 ± 8.76 | $F_{3, 59}$ = 10.58<br>*P* < 0.001 |
| Escape $\Delta\beta$ | N = 32<br>M = 0.03 ± 0.07 | N = 11<br>M = -0.02 ± 0.03 | N = 19<br>M = -0.03 ± 0.05 | $F_{3, 68}$ = 5.33<br>*P* = 0.002 |
| Sustained $\Delta\beta$ | N = 34<br>M = -0.01 ± 0.03 | N = 12<br>M = 0.02 ± 0.02 | N = 19<br>M = 0.02 ± 0.03 | $F_{3, 71}$ = 7.28<br>*P* < 0.001 |
| BISQ: nr wakings per night | N = 40<br>M = 2.81 ± 0.95 | N = 13<br>M = 2.65 ± 1.09 | N = 17<br>M = 2.38 ± 0.86 | $F_{2, 67}$ = 1.21<br>*P* = 0.304 |
| BISC: time to sleep | N = 40<br>M = 21.25 ± 16.52 | N = 13<br>M = 20.00 ± 16.71 | N = 18<br>M = 16.11 ± 5.30 | $F_{2, 68}$ = 0.77<br>*P* = 0.465 |
| BISC: is sleep a problem | N = 40<br>M = 0.35 ± 0.48 | N = 13<br>M = 0.69 ± 0.63 | N = 18<br>M = 0.67 ± 0.59 | $F_{2, 68}$ = 3.21<br>*P* = 0.046 |
| BISQ: how regular is sleep | N = 40<br>M = 1.88 ± 0.65 | N = 13<br>M = 2.31 ± 0.48 | N = 18<br>M = 2.06 ± 0.64 | $F_{2, 68}$ = 2.49<br>*P* = 0.090 |
| BISQ: how easy to wake | N = 40<br>M = 0.88 ± 0.56 | N = 13<br>M = 1.00 ± 0.71 | N = 18<br>M = 1.17 ± 0.79 | $F_{2, 68}$ = 1.26<br>*P* = 0.291 |
| Median night - day sleep duration | N = 39<br>M = 3.38 ± 1.27 | | | |
| Daytime sleep duration | N = 39<br>M = 37.11 ± 6.85 | | | |
| Nighttime sleep duration | N = 39<br>M = 60.85 ± 5.91 | | | |
| Total sleep duration | N = 39<br>M = 97.96 ± 9.68 | | | |
| Sleep duration variability | N = 39<br>M = 2.00 ± 0.55 | | | |
| Nr daytime sleep transitions | N = 39<br>M = 68.31 ± 14.70 | | | |
| Nr nighttime sleep transitions | N = 39<br>M = 56.77 ± 14.66 | | | |
| Total nr sleep transitions | N = 39<br>M = 115.64 ± 22.76 | | | |

$\Delta\beta$ denotes the difference in slopes between blue and red light stimulation: $\Delta\beta = \beta_{red} - \beta_{blue}$. All durations are measured in hours. Sleep transitions include both wake-to-sleep and sleep-to-wake transitions. Shaded cells indicate ANOVAs with significant post-hoc-tests using Bonferroni correction; the row “BISC: is sleep a problem” did not have any significant post-hoc differences between any groups and is therefore white. ± is s.e.m.

**Supplementary Table 2.** Preregistered analyses using linear mixed models and the fitlme function.

| **Wilkinson model** | **Estimate (95% CI)** | **F-test** | **P-value** |
|---|---|---|---|
| Δ***β*** ~ ageInMonths | 0.004 (0.002 - 0.006) | F(1, 58) = 14.489 | 0.000 |
| BISQ_nrWakingsPerNight ~ Δ*β* | -0.972 (-8.798 - 6.854) | F(1, 57) = 0.062 | 0.805 |
| BISQ_timeToSleep ~ Δ*β* | -22.335 (-145.066 - 100.396) | F(1, 58) = 0.133 | 0.717 |
| BISQ_isSleepAProblem ~ Δ*β* | 3.479 (-0.767 - 7.726) | F(1, 58) = 2.690 | 0.106 |
| BISQ_howRegular ~ Δ*β* | 1.129 (-4.026 - 6.285) | F(1, 58) = 0.192 | 0.663 |
| BISQ_howEasyToWake ~ Δ*β* | 0.882 (-4.158 - 5.923) | F(1, 58) = 0.123 | 0.727 |
| daytimeSleepDurationHours ~ Δ*β* | -8.648 (-71.105 - 53.810) | F(1, 28) = 0.080 | 0.779 |
| nighttimeSleepDurationHours ~ Δ*β* | 2.381 (-62.863 - 67.625) | F(1, 28) = 0.006 | 0.941 |
| totalSleepDurationHours ~ Δ*β* | -6.267 (-108.616 - 96.082) | F(1, 28) = 0.016 | 0.901 |
| sleepDurationVariability ~ Δ*β* | 3.099 (-2.397 - 8.596) | F(1, 28) = 1.334 | 0.258 |
| nrDaytimeSleepWakeTransitions ~ Δ*β* | 25.437 (-117.349 - 168.223) | F(1, 28) = 0.133 | 0.718 |
| nrNighttimeSleepWakeTransitions ~ Δ*β* | -75.349 (-234.556 - 83.858) | F(1, 28) = 0.940 | 0.341 |
| nrTotalSleepWakeTransitions ~ Δ*β* | -27.105 (-271.309 - 217.099) | F(1, 28) = 0.052 | 0.822 |

These analyses were preregistered, but in the main text we present data grouped into 2-month-, 6-month- and 10-month-olds to simplify visual presentation and we there present the data using 1-way ANOVAs and post-hoc follow up. All effects are similar regardless of statistical method and does not change any interpretations of the results. Shaded cells indicate significant effects.

**Supplementary figures and legends**

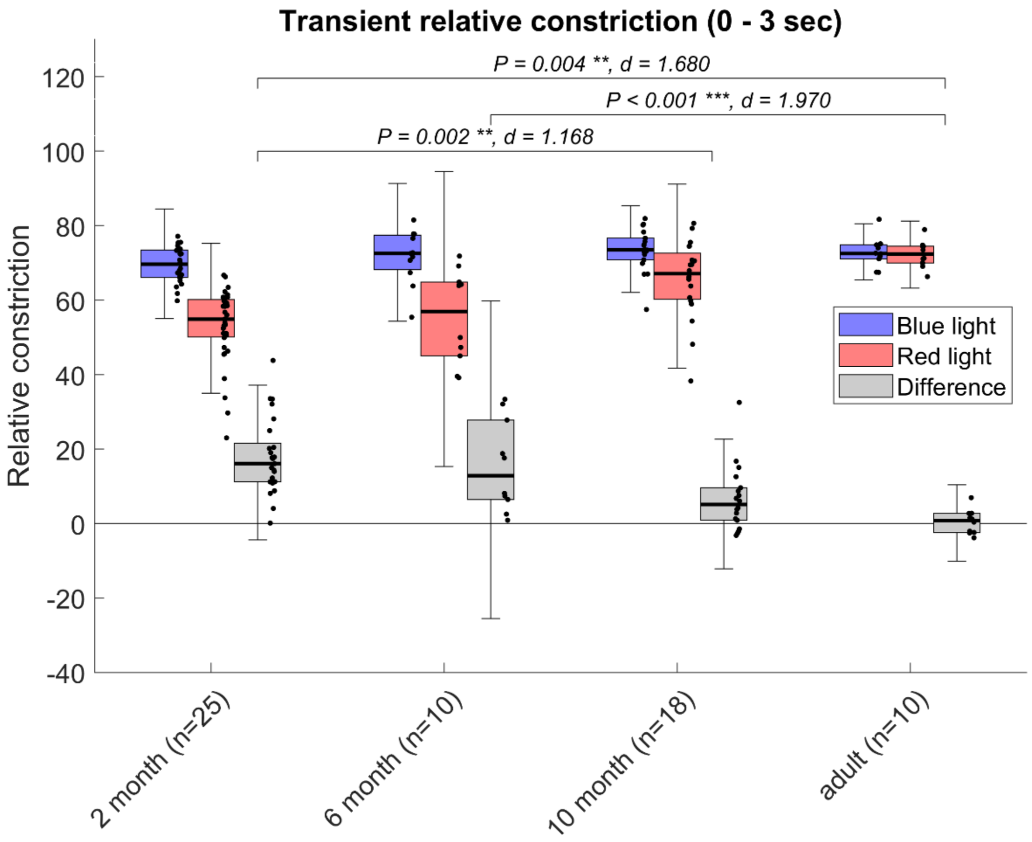


**Supplementary Figure 1.** Post-hoc test of relative constriction after light onset shows decreased differences between blue-red constrictions when comparing the 2-month-olds and the 10-month-olds. Only the difference values between red and blue stimulation were used for statistical analysis. Effect sizes are Cohen's d, which are presented together with P-values above significance brackets. Center line is median; box limits are upper and lower quartiles; whiskers are 1.5x interquartile range; points are individual participant values.

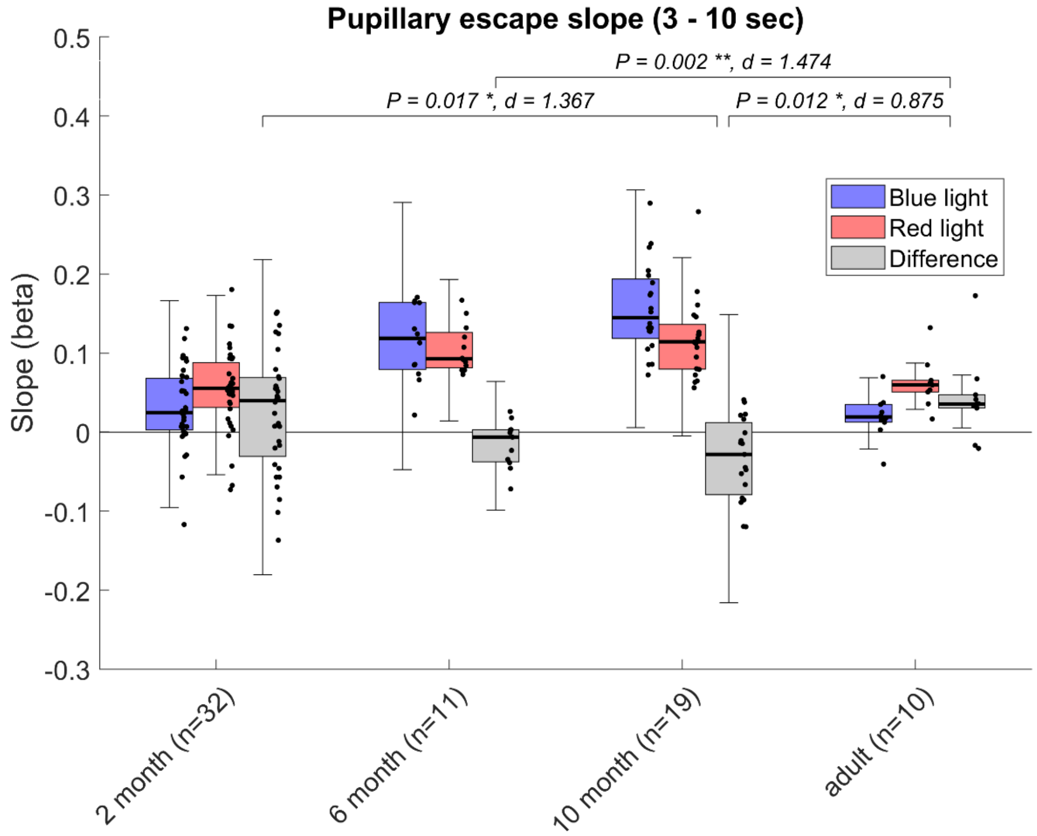


**Supplementary Figure 2.** Post-hoc test of pupillary escape shows a change between 2-month-olds and 10-month-olds, mostly driven by a larger pupillary escape to blue light stimulation across ages. Only the difference values between red and blue light stimulation were used for statistical analysis. Effect sizes are Cohen's d, which are presented together with P-values above significance brackets. Center line is median; box limits are upper and lower quartiles; whiskers are 1.5x interquartile range; points are individual participant values.
